# Red light imaging for programmed cell death visualization and quantification in plant-pathogen interactions

**DOI:** 10.1101/2020.08.14.251009

**Authors:** Sergio Landeo Villanueva, Michele C. Malvestiti, Matthieu H.A.J. Joosten, Wim van Ieperen, Jan A.L. van Kan

## Abstract

Studies on plant-pathogen interactions often involve monitoring disease symptoms or responses of the host plant to pathogen-derived immunogenic patterns, either visually or by staining the plant tissue. Both these methods have limitations with respect to resolution, reproducibility and the ability to quantify the results. In this study we show that red light detection in a multi-purpose fluorescence imaging system that is probably available in many labs can be used to visualize plant tissue undergoing cell death. Red light emission is the result of chlorophyll fluorescence upon thylakoid membrane disassembly during the development of a programmed cell death process. The activation of programmed cell death can occur either during a hypersensitive response to a biotrophic pathogen or an apoptotic cell death triggered by a necrotrophic pathogen. Quantifying the intensity of the red light signal enables to evaluate the magnitude of programmed cell death and provides a non-invasive readout of the plant immune response in a faster and safer manner as compared to chemical staining methodologies previously developed. This application can be implemented to screen for differences in symptom severity in plant-pathogen interactions, and to visualize and quantify in a sensitive and objective manner the intensity of a plant response upon perception of a given immunological pattern. We illustrate the utility and versatility of the method using diverse immunogenic patterns and pathogens.

## Introduction

In natural environments as well as in agricultural systems, plants are constantly threatened by pathogens, including nematodes, oomycetes, fungi, bacteria and viruses. In order to invade, colonize and obtain nutrients from their host, these pathogens must deal with plant defense mechanisms and therefore they have evolved different lifestyles and strategies to overcome the plant immune system and cause disease. Given their sessile nature, plants rely in the first line of their defense on anatomical barriers and preformed phytochemical compounds to prevent pathogen invasion. However, when pathogens can overcome these first passive defense layers, recognition of the invaders by the plant can take place within the host tissue. To perceive danger, plants have evolved a wide range of receptors and their function consists of the recognition of so-called immunogenic patterns (IPs), both in the apoplast and in the cytoplasm (Van der Burgh & Joosten, 2019; Gust et al., 2017). Triggers of plant immunity include damage-associated molecular patterns (DAMPs) (Boller & Felix, 2009), structural components of the invaders (microbe-associated molecular patterns, MAMPs), and pathogen-secreted compounds known as effectors that serve to perturb plant defense responses and promote virulence (Kanyuka & Rudd, 2019). Apoplastic triggers of immunity are referred to as extracellular immunogenic patterns (ExIPs) (Van der Burgh & Joosten, 2019). These are perceived by trans-membrane pattern recognition receptors (PRRs) located on the cell surface, which have been grouped into receptor-like kinases (RLKs, containing a cytoplasmic kinase domain) and receptor-like proteins (RLPs, which lack such a domain) (Gust et al., 2017). By contrast, cytoplasmic danger signals are referred to as intracellular immunogenic patterns (InIPs) and are recognized by the activity of cytoplasmic receptors, usually from the category of nucleotide binding leucine-rich repeat receptors (NB-LRRs) (Van der Burgh & Joosten, 2019). Extra- and intracellular receptors possess highly variable LRR domains, which confer specific binding capacity to a wide array of ExIPs and InIPs (Van der Burgh & Joosten, 2019). Upon perception of immunogenic patterns, PRRs and NB-LRRs activate down-stream signaling pathways leading to the induction of immune responses to block the invader.

Plant immune responses comprise multiple cellular processes, among others, alkalization of the extracellular space (Granado et al., 1995; Jourdan et al., 2009), cell wall reinforcement via callose deposition (Luna Diez et al., 2010; Voigt, 2014), accumulation of reactive oxygen and nitrogen species (Turkan, 2018), an increase in intracellular calcium (Grant et al., 2000), biosynthesis of pathogenesis-related (PR) proteins (Jain & Khurana, 2018) and phytoalexins (Jeandet, 2015) and the release of small RNAs to interfere with the transcriptional machinery of the invading pathogen (Cai et al., 2018). In an attempt to stop pathogen invasion and disease development, the affected plant cells can commit suicide by undergoing programmed cell death (PCD). Even though it represents a decisive and seemingly altruistic response of an individual plant cell to pathogen invasion, PCD can lead to either resistance or to disease development, depending on the lifestyle of the pathogen. In incompatible interactions with biotrophic pathogens, PCD is referred to as the hypersensitive response (HR) and manifests itself as an effective resistance response of the plant upon pathogen recognition. The HR represents a form of autophagic PCD, preventing the spread of biotrophic microbes (Michaeli et al., 2016; Kabbage et al., 2017). For example, race 5 of the biotrophic fungus *Cladosporium fulvum*, which causes leaf mold of tomato following a typical gene-for-gene interaction, cannot cause disease on tomato carrying the gene encoding the RLP Cf-4 (Joosten et al., 1994; Thomas et al., 1997). This PRR recognizes the Avr4 effector protein that is secreted by *C. fulvum* race 5 and mediates the activation of the HR, leading to leaf mold resistance (Thomas et al., 1997). On the other hand, in interactions with necrotrophs, PCD occurs after induction of apoptosis of the affected host cells and becomes visible as a necrotic lesion (Curtis & Wolpert 2004, van Baarlen et al., 2004). In this situation, PCD shows specific apoptosis hallmarks, which include cell shrinkage and membrane blebbing with retention of plasma membrane integrity; loss of mitochondrial membrane potential and release of cytochrome c into the cytoplasm; protease activation; chromatin condensation and nucleosome cleavage; decrease in ATP and increase in reactive oxygen species (ROS; Dickman et al., 2017). Necrotrophic pathogens, such as fungi of the genus *Botrytis*, exploit the plant PCD pathway by actively triggering apoptosis in host cells through the activity of an array of effector compounds, allowing the pathogen to acquire nutrients from the dead plant cells and to colonize the plant tissue (Veloso & van Kan, 2018).

In order to determine the level of resistance or susceptibility of a given plant genotype, assessment of disease symptoms in an infection assay is a key step to evaluate the outcome of an interaction between host and pathogen. Thus, visualization and quantification of PCD over a given period of time are essential to understand how the plant responds upon contact with the pathogen. Furthermore, a visual assessment is often performed to study the effect of individual pathogen IPs (including effectors) on the host, which is done by applying such molecules to a plant tissue, either by agro-infiltration or IP injection. This approach allows to compare plant genotypes for their responses to pathogens or specific IPs and to monitor disease development or the establishment of resistance responses over time.

At present, the most widely used and commonly accepted method to visualize plant cell death is the trypan blue staining (TBS) technique, which is a cell viability test based on the principle that live cells, possessing intact and functional cell membranes, exclude the trypan blue dye whereas dead cells do not. This staining method was first described by Ehrlich (1885) in a study in which trypan blue dye was injected into animal tissue and he observed that the brain tissue showed less staining. Subsequent research showed that when trypan blue was injected directly into the central nervous system instead, the brain tissue would stain equally well, but the stain would not travel to the rest of the body, showing a compartmentalization between the cerebrospinal fluid and the vasculature in the rest of the body (Ehrlich, 1885; Mott, 1913). In the course of the 20^th^ century and until current times, TBS has been successfully implemented in plant biology and phytopathological studies, since it was found that TBS allows to visualize plant vascular tissue (Escamez et al., 2017), dead Arabidopsis (*Arabidopsis thaliana*) cells as a consequence of HR induced by *Pseudomonas syringae* effectors (Johansson et al., 2015), or cell death responses in *Brassica rapa* cotyledons to oomycete elicitins (Takemoto et al., 2005). TBS became consequently useful to assess the extent of plant tissue colonization by microbial pathogens, and for example for detecting the presence of (micro-) lesions in plant-microbe interactions (Vogel et al., 2005). Nevertheless, TBS has four major drawbacks. First of all, it is not understood how the trypan blue dye can pass membranes of dead cells as well as of hyphae of live fungi and oomycetes, whereas the dye is specifically excluded from live plant cells with an intact plasma membrane. Second, the method requires the use of toxic chemicals, such as the dye itself, phenol and chloral hydrate. Third, TBS represents a destructive assay which does not allow to follow disease progression and/or the occurrence of plant cell death responses in the same sample over time, nor does it allow to sample the tissue for RNA, protein or metabolite analyses after it is stained. Finally, the method provides a qualitative test of cell viability, but it does not easily allow to quantify PCD in a standardized way.

We here report a non-destructive, non-toxic and reproducible method to visualize and quantify cell death in green plant tissues. This method can be implemented to study different pathosystems and allows to follow the development of PCD over time. The protocol is based on the detection of light signals emitted by plant cells undergoing cell death. With the aid of a fluorescence imaging system, we observed red light emission by leaf tissue upon infiltration with known PCD-inducing compounds or infection by a pathogen. We provide evidence that the red light signal originates from the increase in chlorophyll fluorescence due to dismantling of chloroplast thylakoid membranes in cells undergoing PCD. We provide several illustrations that this method can be used as a versatile tool for quantitative and non-destructive evaluation of plant responses to microbial pathogens or their IPs.

## Results

This study was initiated after the observation that upon exposure to a green light source, leaf tissue undergoing a typical HR emits light that could be detected using the red light filters of a fluorescence imaging system. Our initial experiments indicated that the intensity of the emitted light positively correlated with the severity of visual signs of cell death. We set out to explore whether this method would be suitable for the development of an alternative, fast, simple and safe, non-destructive tool to visualize cell death in plant tissue. This tool should allow to quantify PCD in a standardized and reproducible manner and to follow the development of PCD over a given time course. Therefore, we first tested whether the areas yielding a light signal that was detected with red light imaging, originated from the affected tissue that in PCD tests is normally stained with trypan blue. *N. benthamiana:Cf-4* leaves were infiltrated with pure Avr4 protein from *C. fulvum* and we compared the visible HR symptoms with the classic TBS and the red light signal on the same leaves (Fig. 1). At 3 days post infiltration, HR symptoms were observed on leaves infiltrated with 50 μM of Avr4 but not 10 μM (Fig. 1, left column). Comparing TBS (Fig. 1, middle column) to the red light signal (Fig. 1, right column), it can be noticed that the areas stained with trypan blue exactly match with the areas from which the red light signal is detected by the imaging system. The signal intensity was clearly stronger at higher Avr4 concentration, and showed a punctate pattern, with the most intense red light emission from areas that displayed the most obvious signs of cell death.

**Fig.1.**
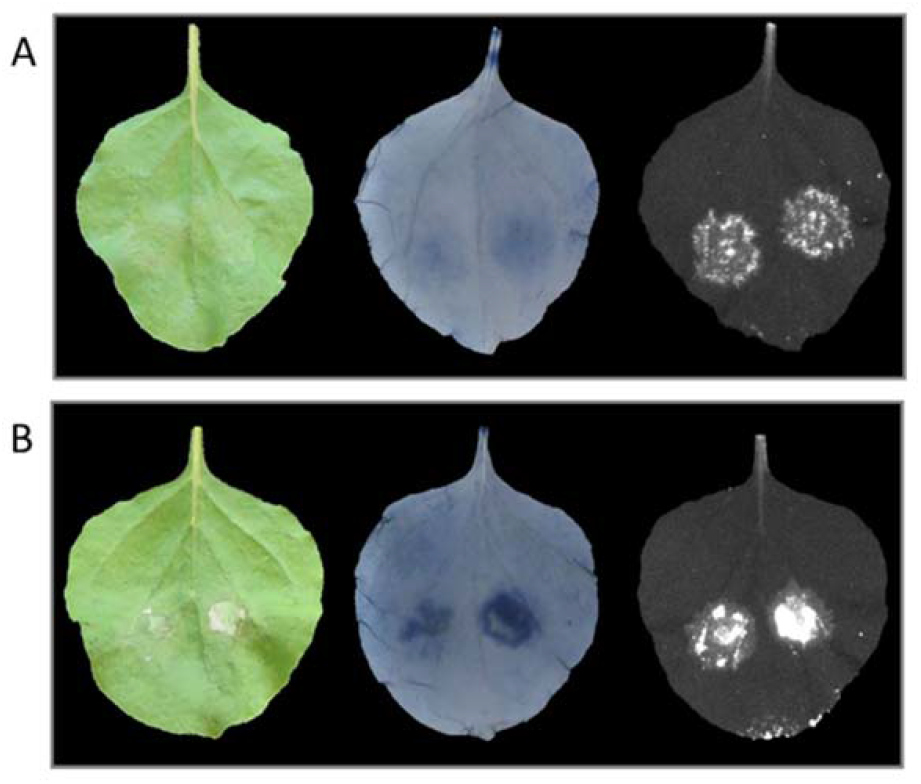
Cell death symptoms caused by Avr4 infiltration (**A**: 10 μM; **B**: 50 μM) in leaves of *N. benthamiana:Cf-4* plants, visualized at 3 dpi with visible light (left column), TBS (middle column) and red light imaging system (right column).

Recently, several studies have used UV transillumination to monitor cell death responses (Guo et al., 2020; Zdrzalek et al., 2020). We compared the use of UV light as a source of excitation for imaging of PCD undergoing tissue to the imaging based on green light excitation described above (Supplementary Fig.S1). When the leaves were excited with UV light, a much stronger background signal was detected in the healthy tissue and in the leaf veins. Such a background signal interferes with the red light-signal that derives from PCD undergoing tissue, thereby rendering the UV imaging system less sensitive for areas yielding lower signal intensities.

Having shown that the necrotic areas exactly match with the areas from which the red light signal originates, we infiltrated different IPs into leaves of *N. benthamiana* to test whether the intensity of the red light signal correlates with the severity of the visible PCD symptoms (Fig. 2 and 3). Upon infiltration of flg22, at 3 dpi no cell death symptoms were observed on the leaves, and also no red light signal was detected (Fig. 2A). On the other hand, a clear red light signal was detected upon Avr4 infiltration in leaves of *N. benthamiana* stably expressing Cf-4, but, apart from some chlorosis, no cell death symptoms were observed (Fig. 2B). In contrast, BcNEP1 infiltration caused the swift development of a clearly visible area of dry necrotic tissue that was matching with the area yielding a strong red light signal (Fig. 2C). In a reactive oxygen species (ROS) burst assay, the incubation of leaf disks of *N. benthamiana* in a solution containing either flg22, Avr4 or BcNEP1, triggered a strong ROS burst (Fig. 2, lower panel).

**Fig.2.**
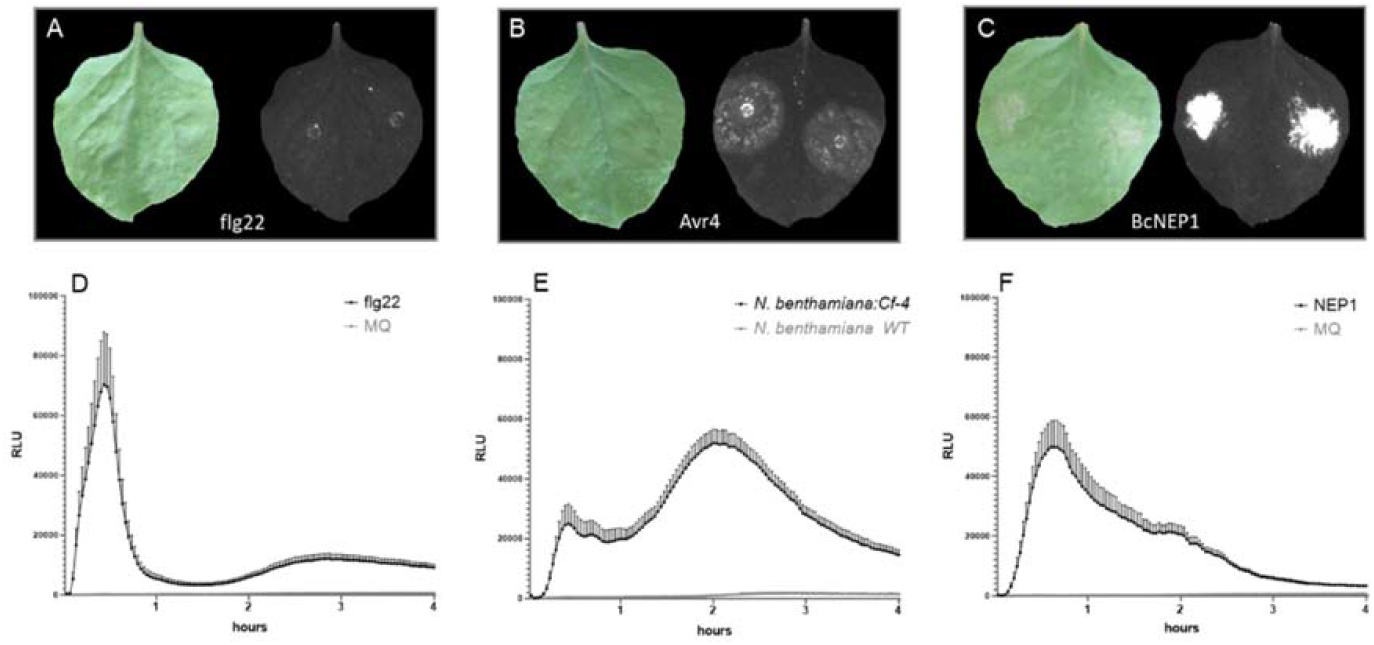
Visible cell death symptoms (left) and detected red light signal (right), upon infiltration of 0.1 μM flg22 (**A**), 0.1 μM Avr4 (**B**) and 0.004 μM BcNEP1 (**C**) at 3 dpi in leaves of *N. benthamiana:Cf-4*. ROS burst triggered by 0.1 μM flg22 and the MQ control in *N. benthamiana* WT (**D**), 0.1 μM Avr4 in *N. benthamiana:Cf-4* and the *N. benthamiana* WT control plant (**E**), and 0.004 μM BcNEP1 and the MQ control in *N. benthamiana* WT (**F**). N=8, error bars show the standard error.

**Fig.3.**
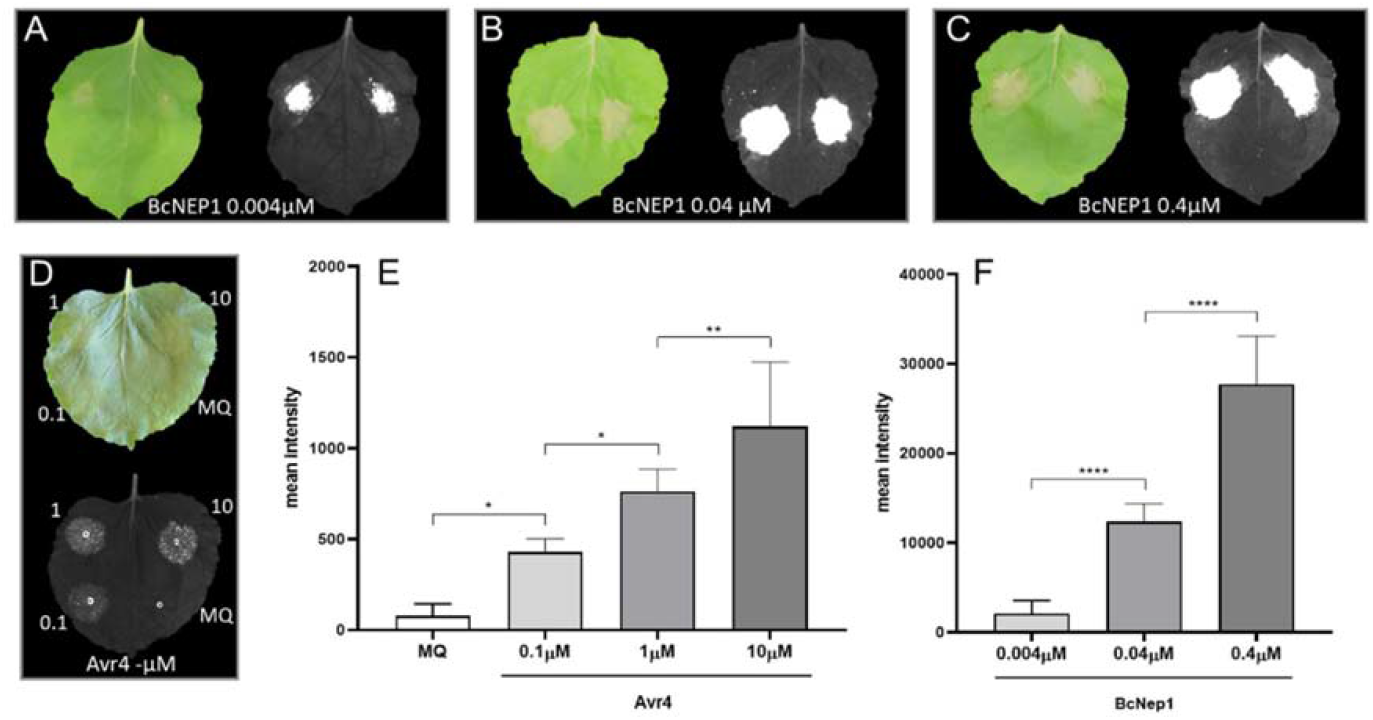
Dose-dependent cell death symptoms caused by BcNEP1 (**A**, **B** and **C**) and Avr4 (**D**) protein infiltration in *N. benthamiana:Cf-4* at 3 dpi, as visualized under visible light and with the red light imaging system. Quantification of red light signal intensities emitted by *N. benthamiana:Cf-4* leaves infiltrated with 0.1, 1 and 10μM Avr4 protein and MQ water as a control (**E**), and with 0.0004, 0.004 and 0.4μM BcNEP1 protein (**F**). Bars represent averages ± standard deviation (n=8). Asterisks indicate significant differences using ANOVA followed by a Tukey test.

To explore whether the imaging system also functions for plants other than *N. benthamiana*, we infiltrated a different IP, namely the *B. cinerea* polygalacturonase 3 (BcPG3) protein, in detached leaves of WT Arabidopsis Col-0 plants and in an *Atsobir1* Arabidopsis mutant (Zhang et al., 2014). As depicted in Supplementary Fig. S2, necrotic tissue in combination with a clear red light signal was observed only in leaves of the WT plants, whereas neither necrotic spots nor a red light signal were detected upon BcPG3 infiltration in the *Atsobir1* mutant. In addition, a specific red light signal was detected in leaves of transgenic tomato seedlings systemically expressing both the *C. fulvum* Avr4 and the Cf-4 resistance protein (Joosten *et al.,* 1994), even before symptoms of necrosis were observed (Supplementary Fig. S3).

Besides the qualitative visualization of PCD in different plant species, this methodology should also allow to quantify the magnitude of PCD by measuring the intensity of the red light signal. To test this application, we infiltrated leaves of *N. benthamiana:Cf-4* with different concentrations of Avr4 protein from *C. fulvum* and BcNep1 protein from *B. cinerea* (Fig. 3). While even the highest concentration (10μM) of Avr4 protein barely induced any sign of PCD as observed by naked eye (Fig. 3D), the imaging system detected increasing red light signal intensities emitted by the infiltrated tissue upon infiltration of higher Avr4 concentrations (Fig. 3D and 3E). By contrast, no red light signal was detected in the mock-infiltrated areas, except at sites where the syringe was applied and pressed on the leaf to infiltrate the MQ, thereby causing some wounding (Fig. 3D, MQ). On the other hand, BcNEP1 infiltration caused visible necrotic spots already at the lowest concentrations used (Fig. 3A), whereby the red light signal intensity measured upon BcNEP1 infiltration was up to 25-fold higher than in Avr4 infiltrated leaves (Fig. 3E and 3F). In addition, it can be noticed that when BcNEP1 was infiltrated at higher concentrations no clear differences in the necrotic tissues could be observed (Fig. 3B and 3C). However, when quantified with the imaging system, increasing red light signal intensities were recorded in leaves infiltrated with increasing BcNEP1 concentrations (Fig. 3F).

Furthermore, to monitor the development of PCD over a given time course, leaves of *N. benthamiana* and Arabidopsis Col-0 were inoculated with a conidial suspension of *B. cinerea*. In both cases, development of necrotic lesions was observed from 2 dpi and the imaging system detected a clear red light signal at the site of the lesion (Fig. 4A and 4B). For *N. benthamiana*, pictures of the same detached leaves were taken at 2, 3 and 4 dpi. This was not possible for Arabidopsis leaves because they were too fragile after detachment, and therefore symptom development was followed upon detaching different leaves at 2 and 3 dpi. Following incubation, we observed that the lesions were expanding and that the red light signal was sometimes detected in irregular shapes, outside of the circular necrotic spot caused by the fungus. The latter light-emitting zones were not visible by the naked eye (Fig. 4A and 4B, yellow arrows). Moreover, the imaging system detected a red light signal for both Arabidopsis and *N. benthamiana*, which was visible as concentric rings of different intensities at the location where the necrotic lesions developed (Fig. 4A and 4B, red arrows).

**Fig.4.**
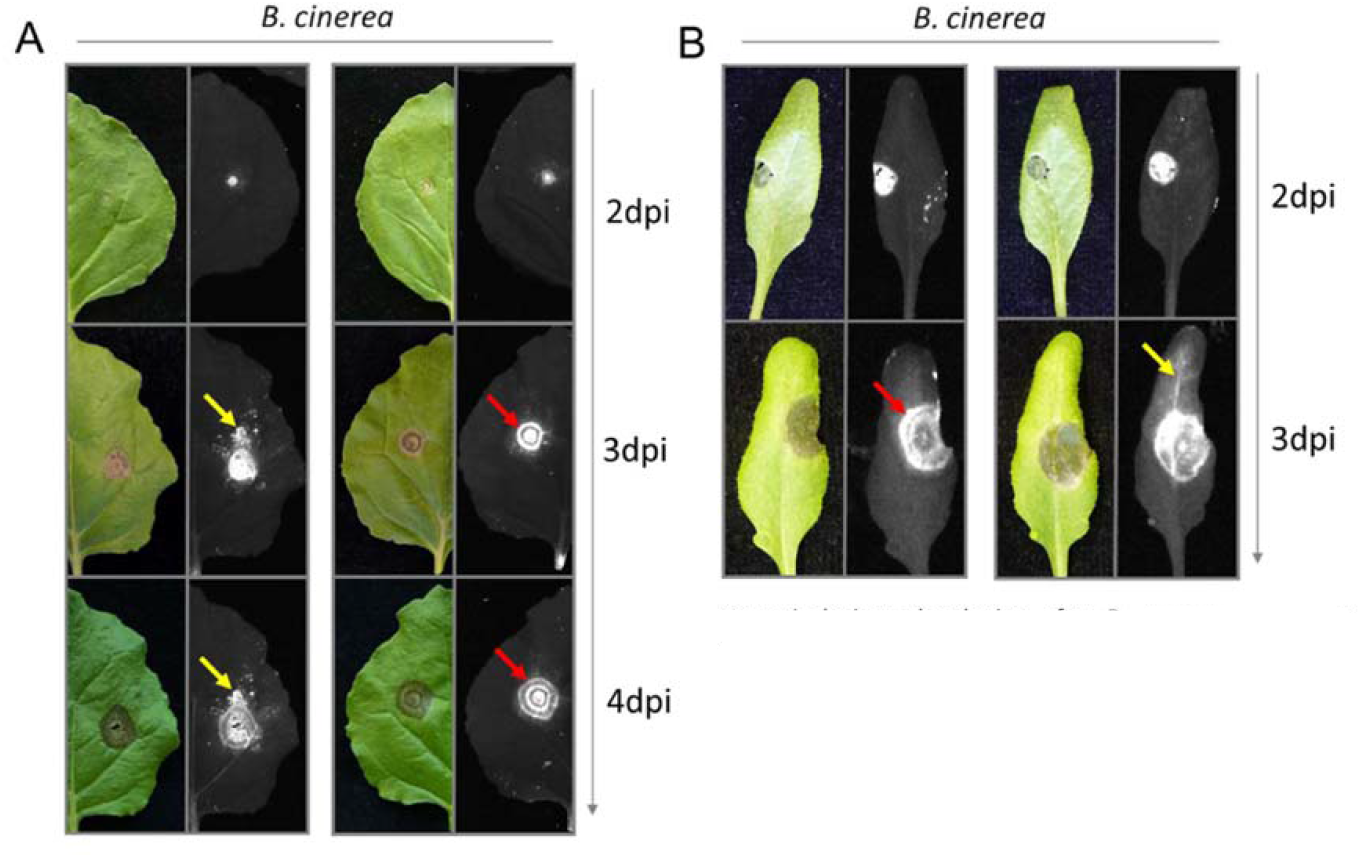
Necrotic lesions developing after *B. cinerea* inoculation on leaves of (**A**) *N. benthamiana* and (**B**) Arabidopsis, visualized at visible light (left) and with the red light imaging system (right). Yellow arrows indicate the red light signal detected in tissue outside the necrotic lesions; red arrows indicate distinct concentric rings with different red light signal intensities within the lesion.

We also followed the development of necrotic lesions on two lily cultivars inoculated with two different *B. elliptica* isolates. Fig. 5 shows the necrotic lesions that were observed at 3 and 4 dpi. In all cases the lesions expanded during the incubation. With both isolates, distinct areas were observed emitting a light signal of different intensities, with a fairly abrupt transition between them (Fig. 5B). The outer area showed a lower red light intensity (Fig. 5B) and this was not visible as necrotic tissue when the picture was taken with visible light. Moreover, for the most aggressive isolate (*Be9401*) in both cultivars the imaging system detected much weaker red light signal in the center of the lesion compared to the concentric rings around the inoculation site, and red light signal was also detected in leaf veins. In contrast, for the less aggressive isolate (*Be9732*), the signal remained intense at the inoculation site, while the lesions were still expanding and no red light signal was detected in the leaf veins. To illustrate the differences in symptom severity and to compare isolate virulence and cultivar susceptibility, the signal intensities over the lesion areas were quantified and the values were plotted on a bar chart (Supplementary Fig. S4).

**Fig.5.**
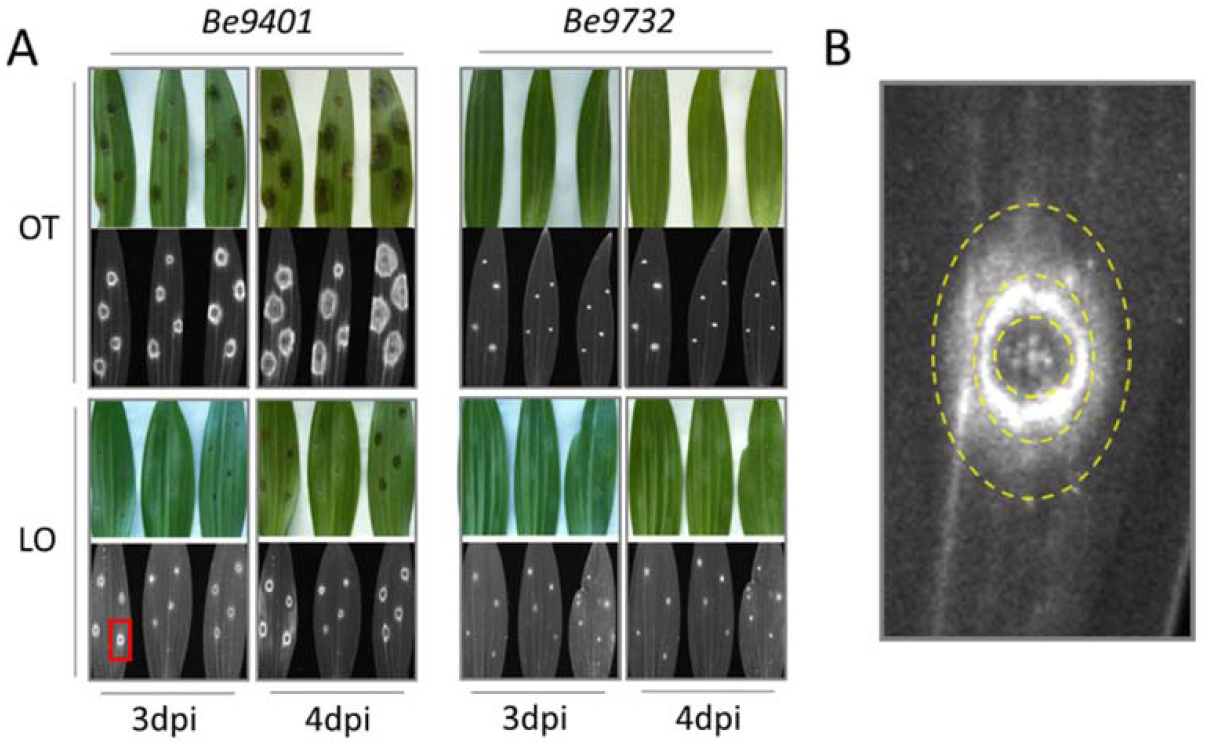
(**A**) Necrotic lesions developing after *Be9401* and *Be9732* inoculation of leaves of lily Cv OT and Cv LO, visualized at visible light (upper) and with the red light imaging system (bottom), at 3 and 4dpi. For each time point and cultivar, three representative leaves are shown. (**B**) Zoom-in on a necrotic lesion observed at 3dpi on Cv LO, highlighted by a red rectangle in (**A**), showing an area with different light signal intensities. Yellow dashed-lines indicate the borders between these areas.

To explain the origin and nature of the observed red light signal in PCD-undergoing tissue, we hypothesized that the signal might derive from the disassembly of thylakoid membranes of the chloroplasts. To test this hypothesis, chloroplasts were isolated from *N. benthamiana* leaves and treated with different concentrations of the membrane-disrupting surfactants Nonidet-P40 and SDS, and the surfactant solutions were also directly infiltrated in detached *N. benthamiana* leaves (Supplementary Fig. S5 and S6). As shown in Fig. 6A and Fig. 6C, red light emission was detected in the chloroplast suspension after treatment with both surfactants, whereas no red light was emitted by untreated chloroplast suspension and in the pure surfactant solutions. The graphs in Fig. 6B and Fig. 6D show that the red light intensities increased when the chloroplasts were treated with a higher concentration of the surfactant, as well as with a longer incubation time with 10% NP40, and with 10% and 20% SDS. Upon direct leaf infiltration with the surfactant solutions, the tissue collapsed in a dose-dependent manner and eventually showed a brownish coloration that resemble the necrotic tissue observed upon infiltration of various IPs, and in the inoculation assays (Supplementary Fig. S3 and S4).

**Fig.6.**
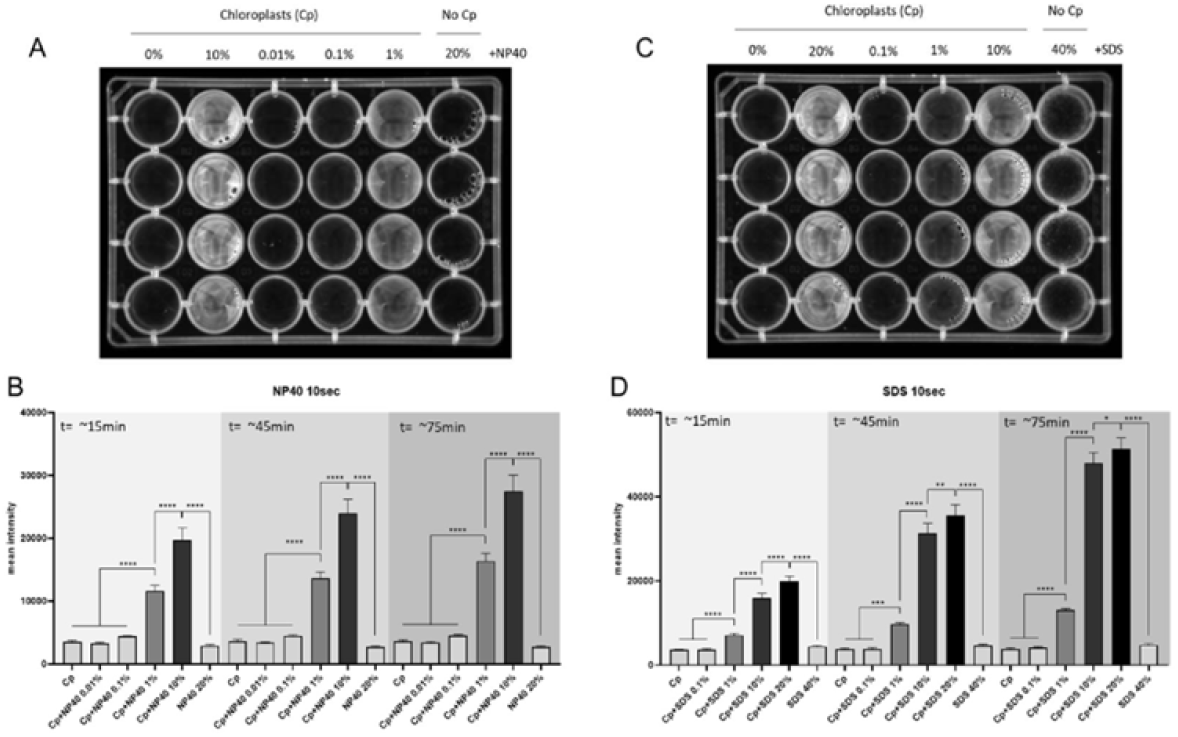
Red light emission in a 24 wells plate containing a chloroplast suspension obtained from *N. benthamiana* leaves, after treatment with respectively NP40 (**A** and **B**) and SDS (**C** and **D**) at different concentrations. Red light signal intensities, determined at three time points after the indicated treatments, were quantified and bars represent averages ± standard deviation (n=4). Asterisks indicate significant differences using ANOVA followed by a Tukey test.

## Discussion

### Advantages and applications of red light imaging for studies in plant pathology

Various imaging systems have been developed to investigate plant defense responses. These include quantification methods for ion flux changes and ROS accumulation (Grant & Loake, 2000), nitric oxide generation (Delledonne et al., 1998) and biophoton generation (Bennett et al., 2005). Despite focusing on plant defense responses, none of these methods allows to investigate PCD as a determinant of the outcome of the interaction in a given pathosystem, which usually requires a destructive staining method, TBS. In this study we describe a simple methodology to visualize and quantify PCD in green plant tissues by using the red light filter of a fluorescence-luminescence imaging system, which was originally purchased for imaging DNA or protein gels, and Western blots. We here show that the system can also be applied for imaging plant cell death during interactions with pathogens or their IPs.

Compared to classical TBS, the red light imaging method shows several advantages. First of all, it is safer since it does not require the use of toxic chemicals for preparing the staining and de-staining solutions, and it generates no chemical waste. The method is fast and simple because it is solely based on the detection of a light signal that is automatically processed by the imaging system. This allows to analyze a large number of samples in short time, which makes it useful for large-scale screenings, with up to hundreds of leaves per day. Importantly, in contrast to TBS, this method is non-destructive, and the same samples can be analyzed at different time points to monitor the progression of PCD or the imaging may assist in selecting different regions in the sample for extraction of RNA, proteins or metabolites. Both in case of leaf infiltration with different IPs and in inoculation assays with pathogens, the method allows to observe the plant response at different time points, thereby enabling to follow the development of symptoms during the interaction over a given time span (Fig. 4 and 5). The red light imaging system allows to visualize and quantify PCD in a standardized manner. Different parameters, such as the exposure time and the focus on particular sites of the sample, can be set in advance and with the aid of software, the red light intensities can be quantified for comparison among different samples. On the other hand, for lesion size measurements, it is possible to quantify the area of the tissue of which the red light signal exceeds background levels. In contrast, in TBS the staining intensity may depend on how long the tissue is treated with the staining and de-staining solutions, thereby only permitting a qualitative visualization of PCD, but hampering a quantitative comparison across multiple samples. It is important to note that the imaging system requires placing the samples on a flat surface. Therefore, it is not possible to acquire images from three dimensional objects such as whole plants, since in that case the device is unable to bring the complete area to be studied in focus. More specialised and expensive equipment would be needed for such purposes.

This simple and non-destructive method allows to open new research aimed at studying the role of PCD in plant-pathogen interactions. PCD symptoms can be detected before becoming visible by naked eye and it might be able to distinguish different stages of PCD depending on light intensities. In necrotic lesions caused by *Botrytis* spp., distinct concentric areas were observed showing a red light signal of different intensity. The absence of a red light signal at the center of the lesion (Fig. 5B), indicates that at that location the light signal emitting compounds have been completely degraded. In some cases, when moving distally from the center of the lesion, the first area surrounding the inoculation site showed a higher signal intensity as compared to the lesion periphery, resulting in the formation of a concentric ring (Fig. 5B). This observation might indicate that PCD is more advanced in the brighter region and only just being induced in the peripheral area. It remains to be established whether higher signal intensities correlate with more advanced stages of PCD. Moreover, by inoculating plant tissue with transgenic pathogen strains expressing GFP, it should be possible to investigate whether host tissue colonization has already occurred in the areas displaying low intensity red light signals, or whether PCD has been induced by effector compounds that are secreted by the pathogen and have moved into the surrounding host tissue.

We tested the versatility of the methodology for investigating PCD development by infiltration of different IPs. Upon flg22 infiltration, we neither observed a red light signal nor development of a necrotic lesion, despite the fact that flg22 recognition took place in the leaf tissue and triggered a ROS burst as previously reported (Fig. 2) (Gómez-Gómez et al., 1999). This observation confirms that PCD activation requires specific signaling mechanisms to be induced and that not all IPs equally trigger PCD (Van der Burgh & Joosten, 2019). There was a dose- and time-dependent cell death response upon infiltration of various IPs at different concentrations (Fig. 1 and 3). Since the symptoms are not always clearly visible, the red light filter of the imaging system provides a more reliable and objective tool to quantitatively determine the contribution of given IPs to PCD induction in the host tissue. Finally, this method will help to investigate the function of pathogen-secreted effectors. For example, by labelling pathogen-secreted effectors with fluorescent tags, it will be possible to study their mobility in the tissue and localization over time during host infection, in conjunction with the host response to these effectors. Additionally, by identifying areas that are in an early stage of PCD, showing low intensity red light signals, it should be possible to specifically dissect and process tissue areas to isolate proteins or RNAs at the appropriate time and location. Possibly, in this way also specific effector targets in the host cells and PCD signaling components can be identified via co-immunoprecipitation assays, as the appropriate tissue can be collected using the imaging system.

### PCD, chlorophyll fluorescence and the red light signal

This study shows that PCD-undergoing tissue emits red light that can be detected by means of the imaging system. We hypothesized that the red light signal derives from increased chlorophyll fluorescence, as a consequence of the disassembly of the thylakoid membranes in the chloroplasts. This hypothesis was tested by treating isolated chloroplasts with membrane-disrupting surfactants (Fig. 6) and by leaf infiltration with such surfactants (Supplementary Figures S5 and S6). The membrane-disrupting properties of the surfactants caused a concentration-dependent emission of the red light signal from isolated chloroplasts and infiltrated leaves. From a photochemical point of view, this phenomenon can be explained by the abortion of photochemical activity, due to the dismantling of the chloroplast thylakoid membranes in cells undergoing PCD, resulting in an increase in chlorophyll fluorescence. In photochemically active plant cells, chlorophyll molecules are tightly packed together in the light harvesting complexes (LHCs) that are located at the thylakoid membranes of the chloroplasts. Chlorophyll fluorescence is quenched by photochemical activity when LHC-photo-system II (PSII) complexes are assembled in thylakoids. Chlorophylls have two distinct absorption wavelength peaks in the visible light spectrum, the highest is found in the blue spectrum at ca 450 nm and the lowest at 650 nm in the red spectrum. Moreover, chlorophylls possess a fluorescent emission peak in the red spectrum at ca 680 nm. During PCD or upon perturbation of thylakoid membrane stability, LHC-PSII super complexes are disassembled from chloroplast thylakoid membranes. This causes cessation of the electron flow required for photochemical activity because the connection between PSII and PSI via plastoquinone, cytochrome-b6f and plastocyanin is lost. As consequence, light excited chlorophyll molecules will dissipate photon energy through heat and fluorescence (Baker, 2008).

### Conclusion

In this study we show that red light fluorescence detection by a multi-purpose imaging system that is available in many labs can be used to visualize plant tissue undergoing cell death. Quantifying the intensity of the red light signal enables to evaluate the magnitude of cell death, and thereby provides a non-invasive readout of the plant immune response in a faster and safer manner, as compared to chemical staining methodologies previously developed. This method can be exploited to screen for differences in symptom development in plant-pathogen interactions, and to visualize and quantify in a sensitive and objective manner the intensity of a plant response upon perception of a given immunogenic pattern.

## Materials & Methods

### Plant material and growth conditions

Transgenic *Nicotiana benthamiana* plants stably expressing Cf-4 (*N. benthamiana::Cf-4*) (Gabriëls et al., 2006) and wild type (WT) *N. benthamiana* plants were grown under 16h light at 24°C and 8h darkness at 22°C, at a relative humidity of 75%.

Arabidopsis ecotype Columbia-0 (Col-0) and the *Atsobir1* mutant line in the Col-0 background (Gao et al., 2009), were grown under 12h light at 21°C, and 12h darkness at 19°C, at a relative humidity of 75%.

*Lilium* spp. cultivars OT and LO were planted as bulbs and grown in the greenhouse under natural day/night light regime and temperatures. Mature leaves of two months old plants were used for the inoculation assays.

### Red light imaging and signal quantification

Treated plant material was placed in a ChemiDoc MP imaging system, model Universal Hood III (Bio-Rad) and images were acquired upon excitation by a light source in the green VIS spectrum (Green LED Module Kit #1708284) or in the UV spectrum (UV-B Transillumination #1001361) and, with filters capturing the light emitted in red VIS spectrum (filter 605/50). The exposure time was adjusted to avoid image saturation. For comparison purposes, the shown images were auto scaled according to the treatment that yielded the lowest signal intensity.

For the IP infiltration assays, the mean signal intensity was calculated using Image Lab^Tm^ software (version 6.0.1), by manually selecting the treated areas and subtracting the background signal intensity value. For the disease assay in *Lilium*, the spots yielding a light signal were manually highlighted and the signal was calculated as the intensity over the selected area among 12 inoculations sites on three leaves per plant cultivar and per *Botrytis elliptica* isolate at 2, 3 and 4 dpi.

### Statistical analysis

GraphPad Prism^Tm^ (version 8.4.0) was used to perform an analysis of variance (ANOVA) with a post-hoc Tukey’s test on the IP infiltration experiments, chloroplast treatment with surfactants and the *Botrytis* disease assays in *Lilium*.

### Immunogenic patterns (IP)-triggered cell death assays

NEP1 protein from *B. cinerea* (BcNEP1) was produced in *Pichia pastoris* (Schouten et al., 2007) and dissolved in 10mM phosphate buffer, pH = 5.5. The peptide flg22 (EZBiolab) and the protein Avr4 (produced in *Pichia pastoris;* van den Burg et al., 2001) from *C. fulvum* were dissolved in water. Protein infiltration was carried out in the first fully expanded leaf of 5-6 weeks old *N. benthamiana::Cf-4* and WT plants, and the cell death response was scored at 3dpi. The infiltrated areas were photographed, and the average red light signal intensities were calculated.

*B. cinerea* polygalacturonase 3 (BcPG3) protein produced in *Pichia pastoris* (Zhang et al., 2014), was dissolved in 10mM sodium acetate buffer pH = 4.2 and infiltrated at a concentration of 1.5 μM in fully expanded rosette leaves of 6-7 weeks old Arabidopsis plants. The necrotic areas were photographed and analyzed by red light imaging at 8 dpi. Systemic cell death triggered in Avr4- and Cf-4-expressing tomato seedlings

Tomato (*Solanum lycopersicum*) seedlings stably expressing both Avr4 and Cf-4, were generated by crossing transgenic tomato cv Moneymaker (MM) expressing *C. fulvum* effector Avr4 to MM-Cf-4 plants (Stulemijer et al., 2007; Etalo et al., 2013). After sowing, plants were grown for 4 weeks at 33°C and 100% relative humidity (RH), to suppress systemic PCD. Synchronized systemic PCD was eventually induced by transferring the seedlings to 20°C and 70% RH. First fully developed tomato leaves were harvested at 4 hours after transfer and were photographed.

### Trypan blue staining

*N. benthamiana* leaves infiltrated with BcNEP1 protein were harvested at 3dpi and boiled for 2-5 minutes in staining solution, consisting of 16.6% (m/v) of trypan blue in an ethanol/lactic acid/phenol/water (3:1:1:1) mixture. After this, the leaves were de-stained with chloral hydrate and photographed.

### ROS burst measurements

For each treatment, we used eight leaf discs (5 mm) punched from the first fully expanded leaf of 5-6 weeks old *N. benthamiana:Cf-4* plants. The discs were floated on 50μL of Milli Q water (MQ) in a 96-well plate and kept in the dark at room temperature for 5 hours. The MQ was then removed using tissue paper, replaced by 50μL of MQ and kept in the dark for 1 hour at room temperature. After this, 50μL of solution containing the IPs, 50μM of luminol L-012 (FUJIFILM, Osaka, Japan) and 10μg of horseradish peroxidase were added to each well. The luminescence values were measured using a CLARIOstar plate reader (BMG LABTECH Ortenberg, Germany) over a period of 4 hours.

### Disease assays with *Botrytis* spp

Leaves of wild type Arabidopsis Col-0 and *N. benthamiana* plants were inoculated with conidia of *B. cinerea* (isolate B05.10). The fungus was grown on Malt Extract Agar (50g/L, Oxoid) and sporulation was induced by illumination with UV-A lamps. After harvesting, the concentration of the conidia was adjusted to 10^6/mL, using liquid potato dextrose broth (PDB, Oxoid) at 24g/L PDB, and inoculated in 2μL droplets on the adaxial side of the leaves. Mock inoculation was carried out by applying 2μL droplets of PDB solution. Inoculated plants were then incubated in moist plastic boxes and were kept under the same light conditions under which the plants were grown before. Inoculated leaves were detached at 2 days post inoculation (dpi), photographed, and analyzed by red light imaging. The leaves were then stuck into wet floral foam and returned to the plastic boxes to allow disease progression to take place. This procedure was repeated at 3 and 4 dpi to visualize symptom development. Given the fragility of inoculated *A. thaliana* leaves, it was only possible to record symptom development for *N. benthamiana* leaves.

Detached leaves from two different cultivars of *Lilium* spp. (OT and LO), were abaxially inoculated with 2μL droplets of *B. elliptica* conidia from isolates *Be9401* and *Be9732*, respectively (10^5 conidia/mL in 12g/L PDB). Mock inoculation was carried out with 2μL droplets of PDB solution. Inoculated leaves were incubated in moist plastic boxes and pictures of red light imaging were taken at 3 and 4dpi, with 2 seconds of exposure time. The signal intensities were analyzed in a manually drawn area, encompassing the necrotic lesion for all the cultivar-pathogen combinations. An area of variable size was selected in each control leaf for subtraction of the background signal of healthy tissue.

### Thylakoid disassembly test

Intact chloroplasts were isolated from *N. benthamiana* by stirring cut leaf pieces and following the protocol as described by Ho & Theg (2016). After ultracentrifugation, the percoll layer containing the chloroplasts was transferred to a 24 wells plate with 150uL aliquots per well. Two surfactants with membranolytic activity were added to each well to perturb thylakoid membrane stability. The surfactants were used at final concentrations of 0.1, 1 and 10% for Sodium Dodecyl Sulfate (SDS; m/v=g/100mL) and 0.01, 0.1 and 1% for Nonidet-P40 (NP40; m/v=g/100mL). 150uL of the respective surfactant at the different concentrations were added to the wells containing the chloroplast suspension. Untreated chloroplast suspension and pure surfactants at the highest concentration were used as controls. After gently mixing and 20 min of incubation in the dark, the microtiter plates were placed in the imaging system to detect the light signal using red light imaging, at an exposure time of 10 seconds.

## Acknowledgements

The research of SLV is funded by the Peruvian Council for Science, Technology and Technological Innovation (CONCYTEC) and its executive unit FONDECYT. The research of MCM is funded by NWO-Science domain (NWO-ENW), project GSGT.GSGT.2018.008.

## Author contribution list

SLV and MCM performed the experiments, made figures and graphs, and wrote the draft manuscript; MHAJJ and JALvK supervised the study, designed and finalised the manuscript; WvI interpreted the chlorophyll fluorescence data and participated in the writing.

## Supplementary Figure legends

Fig. S1 Cell death symptoms caused by Avr4 infiltration (**A**: 10 μM; **B**: 50 μM) in leaves of *N. benthamiana:Cf-4* plants, visualized at 3 dpi with visible light (left column), TBS (middle-left column) and red light imaging system using green (middle-right column) and UV light (right column) as excitation wave length.

Fig. S2 Cell death symptoms monitored at 8 dpi, caused by BcPG3 protein infiltration in an Arabidopsis WT Col-0 plant (left panels) and in a *Atsobir1* deletion mutant (right panels). Each leaf has been visualized under visible light with visible light (left) and with the red light imaging system (right).

Fig. S3 Red light signal detected in leaves of Avr4/Cf-4 tomato seedlings (**A**) and in control leaves of only Cf-4-expressing tomato seedlings (**B**), harvested 4 hours after changing of growth conditions to a permissive temperature (see materials and methods).

Fig. S4 Red light signal intensity over the lesion areas measured at 2, 3 and 4 dpi from necrotic lesions developed on lily OT and LO inoculated with *Be9401* and *Be9732*. Bars represent averages ± standard deviation (n=12). Asterisk indicates significant differences using ANOVA followed by Tukey.

Fig. S5 (**A**) Leaf response of *N. benthamiana* infiltrated with NP40 at the indicated concentrations, visualized under visible light (upper panel) and with the red light imaging system (lower panel), at 20 min after infiltration of the surfactant. (**B**) Red light signal intensity values measured at 20 min after infiltration with NP40 at the indicated concentrations. Bars represent averages ± standard deviation (n=3). Asterisks indicate significant differences using ANOVA followed by a Tukey test.

Fig. S6 (**A**) Leaf response of *N. benthamiana* infiltrated with SDS at the indicated concentrations, visualized under visible light (upper panel) and with the red light imaging system (lower panel), at 20 min after infiltration of the surfactant. (**B**) Red light signal intensity values measured at 20 min after infiltration with SDS at the indicated concentrations. Bars represent averages ± standard deviation (n=4). Asterisks indicate significant differences using ANOVA followed by a Tukey test.

